# Challenges of Dry Sanitization to Control *Salmonella* Dry Surface Biofilms

**DOI:** 10.64898/2026.08.06.743265

**Authors:** Vinícius S. A. Vaz, Jéssica de A.F.F. Finger, Raul F. Pereira, Erika S. Silva, Rafael P. Maia, Jean-Yves Maillard, Maristela S. Nascimento

**Affiliations:** Departamento de Engenharia e Tecnologia de Alimentos, Faculdade de Engenharia de Alimentos, Universidade Estadual de Campinas, Unicamp, Campinas, SP, Brasil; Departamento de Estatística, Instituto de Matemática, Estatística e Computação Científica, Universidade Estadual de Campinas, Unicamp, Campinas, SP, Brasil; School of Pharmacy and Pharmaceutical Sciences, Cardiff University, Cardiff, UK

**Keywords:** dry sanitizer, dry surface biofilm, low moisture food, food hygiene

## Abstract

*Salmonella* is a pathogen linked to foodborne outbreaks, including low-moisture foods. Its ability to resist desiccation can contribute to the formation of dry surface biofilms (DSB). This study evaluated the impact of 3 DSB formation protocols (P1=48-h hydrated phase/48-h dry phase, P2=24-h/120-h and P3=8-h/48-h) on the resistance of *Salmonella* DSB to 70% alcohol, a commercial product (based on 0.015% quaternary ammonium and 25% isopropyl alcohol), gaseous ozone (45 ppm), hot air (90 °C) and UV-C light (254 nm). The type of DSB protocol impacted the efficacy of the sanitizers (p < 0.05). The biofilm with the shortest hydration phase showed the greatest susceptibility; three out of the five sanitizers evaluated (70% alcohol, commercial product, and UV-C) promoted significant reductions in P3, with counts below the detection limit (0.8 log CFU/cm²) after 5 to 15 min exposure. Regarding protocols P1 and P2, in general, the best performance was from UV-C, especially against DSB on polypropylene, where it achieved reductions of 1.3 log CFU/cm² for P1 and 2.9 log CFU/cm² for P2 after 15 to 30 min of exposure. In contrast, hot air and ozone showed less effectiveness, with reductions ≤1.2 log CFU/cm². In most scenarios, confocal microscopy images corroborated the plate count results (log CFU/cm²). In summary, our data indicates limited action of dry sanitizers on *Salmonella* DSB, requiring validation and optimization of sanitization processes to ensure the microbiological safety of low-moisture products.

## 1. Introduction

*Salmonella* is one of the leading pathogens responsible for outbreaks of foodborne illness (CDC, 2025), with a significant increase in cases associated with low-moisture foods (LMF), such as peanut butter, breakfast cereals, and chocolate (FAO & WHO, 2022; Liu et al., 2022). It was estimated that this microorganism causes approximately 1.35 million infections per year in the United States (CDC, 2025). Its persistence in LMF production environments is associated with its ability to survive under low-moisture conditions and the formation of biofilms (Ivers et al., 2024).

Biofilms are structured microbial communities attached to biotic or abiotic surfaces, often surrounded by a matrix of extracellular polymeric substances (EPS) (Karygianni et al., 2020; Sadiq et al., 2023; Sauer et al., 2022). This structure provides protection against physical and chemical agents, making it difficult to remove microorganisms during cleaning and disinfection processes (Wang et al., 2025). In industrial environments, the presence of biofilms compromises food safety and can cause operational failures, such as cross-contamination, equipment corrosion, and reduced thermal efficiency due to decreased heat transfer (Milton et al., 2023; Olanbiwoninu & Popoola, 2023)

In LMF processing environments, pathogens such as *Salmonella* can form dry biofilms (DSB), which are more difficult to identify and inactivate compared to hydrated-surface biofilms (HSB) (Ledwoch et al., 2022; Lin et al., 2024; Vaz et al., 2026). These biofilms can form directly at the surface-air interface in environments with low water availability, likely as a result of hydration-desiccation phases (Alonso et al., 2023) or through the accumulation of microorganisms on surfaces (Zhong et al., 2009). Recent studies demonstrated significant morphological, structural, and functional differences between DSB and hydrated surface biofilms (HSB). For example, Lin et al. (2024) observed that S. Typhimurium DSB exhibited a more uniform distribution and greater resistance to sanitization, possibly due to layered structural protection—described as a “sandwich” arrangement, in which cells are encapsulated and compressed between the extracellular matrix and dry residues present on the surface. Furthermore, transcriptomic analyses revealed high functional heterogeneity among DSB cells, with over 60% of cells in a metabolically inactive state, while a fraction-maintained antioxidant and virulence activity, posing a significant risk to the safety of LMF (Lin et al., 2024).

Despite the importance of sanitization as an essential step in food industry hygiene programs, effective control of DSB remains a significant challenge, especially in industries seeking to minimize water use to preserve product stability and prevent microbial growth (Baker et al., 2025; Codex Alimentarius, 2018; Lin et al., 2024; Maillard & Centeleghe, 2023; Nerney et al., 2025). In this context, interest in dry sanitization methods is growing. Among the main dry methods, the following stand out: alcohol-based solutions, which promote protein denaturation, cytoplasmic membrane rupture, and cell lysis (Moerman & Mager, 2016; Otto et al., 2011); hot air, which causes protein denaturation, oxidative damage, and cellular dehydration (Prestes et al., 2024); UV-C light, which exerts photothermal and photochemical effects on DNA and membranes (Otto et al., 2011); and gaseous ozone, which has high oxidative potential, reacting with proteins, enzymes, and nucleic acids (Galié et al., 2018). Recent studies have been exploring the efficacy of these methods against hydrated biofilms of different pathogens. Harada and Nascimento (2021a) evaluated treatments with UV-C, dry heat, gaseous ozone, 70% ethanol, and a commercial sanitizer on *Bacillus cereus* hydrated biofilms formed on stainless steel (SS) and polypropylene (PP). The results indicated that UV-C light on both surfaces and ozone on PP showed reductions of approximately 2 log CFU/cm² after 30 min of exposure, while 70% ethanol and the commercial sanitizer showed the lowest reductions on both surfaces. von Hertwig et al. (2023) evaluated the efficacy of UV-C light, hot air, 70% ethanol, and a commercial isopropyl alcohol/QCA-based product on *Salmonella* HSB on SS and PP. UV-C light was the most effective method on PP (reduction of 3.2 to 4.2 log CFU/cm² after 30 min of exposure) and hot air on SS (reduction of 2.2 to 3.3 log CFU/cm²). Although these methods achieved the greatest reductions, the results indicate that efficacy against Salmonella biofilms remains limited (von Hertwig et al., 2023). Despite these advances, studies evaluating the use of physical agents, such as hot air, UV-C light, and gaseous ozone, for the control of Salmonella WBS remain scarce (Alonzo et al., 2018; von Hertwig et al., 2023), and there are no studies on the action of these sanitizers on *Salmonella* DSB. In this context, the present study aimed to evaluate the efficacy of different dry sanitization methods on three types of Salmonella DSB formed on stainless steel (SS) and polypropylene (PP) coupons.

## 2. Materials and methods

### 2.1 Bacterial strains and inocula preparation

Four *Salmonella* strains were used in this study: *S.* Muenster (P03.2 FEA), *S.* Javiana (P06.1 FEA), *S.* Oranienburg (P07.1 FEA), and *S.* Miami (P10.5 FEA). These strains were isolated from the peanut production chain in Brazil (Nascimento et al., 2018) and were able to form HSB, as reported by von Hertwig et al. (2022). The strains were stored at -80 °C until use. For reactivation, a glass bead containing each strain was transferred to 5 mL of Brain Heart Infusion broth (BHI; Difco, Sparks, MD, USA) and incubated at 37 °C for 18-20 h. Subsequently, cultures were streaked onto Trypticase Soy Agar (TSA; Difco) slants, incubated under the same conditions, and stored at 4 °C for further use. At the beginning of the experiment, each strain was cultured twice in BHI at 37 °C for 18-20 h, followed by streaking on TSA and incubated under the same conditions. A loopful of each strain was then transferred to tubes containing 0.85% saline solution, and the turbidity was adjusted to 0.5 on the McFarland scale (Densimat, bioMérieux, France). Decimal dilutions were prepared in 0.1% peptone water (Difco) and subsequently inoculated into Tryptic Soy Broth (TSB; Difco) to obtain a final concentration of approximately 6 log CFU/mL.

### 2.2 Preparation of the coupon

Coupons made of 304 L stainless steel (SS) and polypropylene (PP), with dimensions of 2.0 x 5.0 x 0.1 cm and surface roughness < 0.5 µm, were used. Before each experiment, the coupons were prepared according to Rosado (2009). First, they were placed in an ultrasonic bath (Ultronique, Brazil) for 15 min at 40 kHz. Then, the coupons were immersed in an anionic surfactant detergent solution, manually brushed, and rinsed with distilled water. Afterward, they were immersed in 70% ethanol for 2 h and subsequently rinsed again with distilled water. Finally, the coupons were sterilized at 121 °C for 30 min.

### 2.3 Dry Surface Biofilm (DSB) formation

The methodology used to form the DSB was adapted from Ledwoch et al. (2019). Three protocol conditions were tested to form the DSB: P1 (48 h wet phase + 48 h dry phase), P2 (24 h hydrated phase + 120 h dry phase), and P3 (8 h hydrated phase + 48 h dry phase). Each DSB protocol consisted of two complete cycles of the respective wet and dry phases to ensure biofilm development and experimental reproducibility (Figure 1).

**Figure 1.**
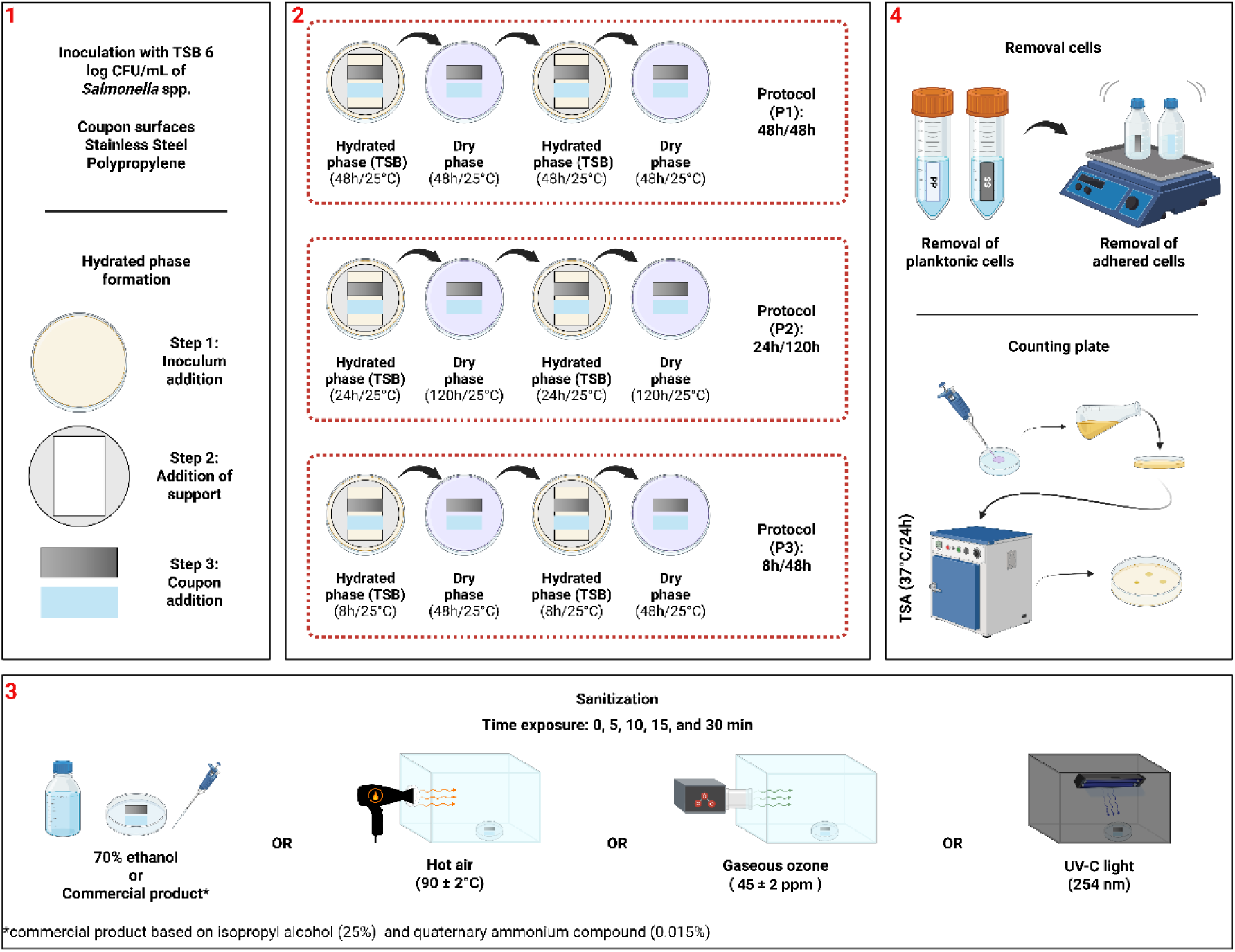
1. Materials; 2. Dry surface biofilm (DSB) protocols; 3. Sanitizer treatments; 4. *Salmonella* determination.

For the DSB formation, SS and PP coupons were placed horizontally over a PP support with a central opening (67 × 43 mm), positioned inside sterile Petri dishes (90 × 15 mm). This setup ensured that only the lower face of each coupon was exposed to the culture medium during the hydrated phase. In this step, 12 mL of TSB, inoculated with a suspension of *Salmonella* strains, was carefully added to the plate to ensure consistent contact with the coupon surface. The plates were then incubated at 25 °C for 8, 24, or 48 h, according to the assigned hydrated phase duration of each protocol. At the end of the hydrated phase, the culture medium was removed using a pipette to avoid disturbing the developing biofilm. The coupons were transferred to new sterile Petri dishes and incubated at 25 °C for the dry phase — 48 h for P1 and P3, and 120 h for P2. This entire process, comprising the wet and dry phases, was repeated once more for each condition, completing two full cycles per protocol. These conditions were designed to simulate varying levels of water stress commonly encountered in LMF environments.

### 2.4 Dry sanitization

At the end of the second dry phase (DSB endpoint), the DSB coupons were transferred to sterile Petri dishes with the biofilm-facing upward and subjected to the five sanitization treatments for 0 (positive control), 5, 10, 15 and 30 min. Coupons without biofilm were analyzed as negative control (Figure 1). All tests were performed three times, as follows:

**I. 70% Ethanol:** The coupons were placed in sterile Petri dishes, and covered with 400 µL of a 70% (v/v) ethanol solution. After the designated exposure time, the coupons were immersed in a neutralizing solution composed of Letheen broth (Acumedia, MI, USA) supplemented with Tween 80 (Sigma Aldrich, Germany) for 5 min (British Standards Institution, 2023). All procedures were carried out under a laminar flow cabinet.
**II. Commercial product:** A commercial product containing 0.015% didecyl dimethyl ammonium chloride (QAC) and 25% isopropyl alcohol was used, following the same procedure described for the 70% ethanol treatment.
**III. Hot air:** The coupons were placed in sterile Petri dishes and transferred to a glass box (30 × 15 × 19 cm). Dry heat was applied using a continuous stream of blown hot air (90 ± 2 °C) generated by a heat blower (Taiff, Brazil). The air temperature was monitored with a thermocouple (PP100, Testo 112, Germany) (Andrade, 2008; Harada & Nascimento, 2021a,b).
**IV. UV-C light:** Before treatment, the UV-C was pre-operated for 30 min to stabilize the UV-C emission. Then the coupons were placed in sterile Petri dishes and positioned inside a dark box (30 × 15 × 15 cm) equipped with an 8 W UV-C lamp (254 nm, 6.5 mW/cm², OSRAM, GmbH, Italy) at a distance of 15 cm (adapted from Sommers et al., 2010). The radiation intensity was monitored using a radiometer (Maestro, Gentec, Canada) (Harada & Nascimento, 2021a).
**V. Ozone:** Sterile Petri dishes containing the coupons were placed inside a hermetic acrylic box (30 × 20 × 20 cm). Gaseous ozone was generated by an ozone generator (Ozoxi, Brazil) connected to a chamber, which was also equipped with an ozone destructor (Ozoxi, Brazil). Ozone concentration was maintained at 45 ± 2 ppm and monitored using an ozone analyzer (UV-100, EcoSensors, USA), following the methodology adapted from Nicholas et al. (2013). Temperature (∼25 °C) and relative humidity (∼50%) were monitored using a sensor (AM2302/DHP22, Aosong, China) (Harada & Nascimento, 2021a)

### 2.5 DSB Quantification

After the exposure time to each sanitizer, the coupons were gently immersed in 15 mL of 0.85% saline solution for 5 min to remove planktonic cells. The coupons that had previously undergone the neutralization step were not immersed in saline solution, as the neutralization step already served as a rinse to remove planktonic cells. Subsequently, they were transferred to a sterile glass bottle containing 50 mL of 0.85% saline solution and 2 g of glass beads. The bottles were shaken (LabLine Orbit Environ – Shaker 3527, LabLine Instruments, IL, USA) at maximum speed for 2 min to release the adherent cells. Decimal dilutions were prepared and plated on TSA, followed by incubation at 37 °C for 24h. The results were expressed as log CFU/cm²

### 2.6 Confocal Laser Scanning Microscopy (CLSM)

CLSM analysis was carried out in SS and PP coupons after 0 (control sample) and 30 min exposure of each sanitization method. The coupons were stained with a 1:1000 dilution of each dye from the FilmTracer™ LIVE/DEAD® Biofilm Viability Kit (Invitrogen, Eugene, OR, USA), which contains the nucleic acid fluorophores SYTO® 9, used to label live cells, and propidium iodide, which labels dead cells. After 10 min of contact with the staining solution, the coupons were washed with PBS and stored in the dark until analysis. Fluorescence images were acquired using a Zeiss LSM780-NLO confocal microscope, coupled to the Axio Observer Z.1 platform (Carl Zeiss AG, Germany), equipped with an EC Plan-Neofluar 20×/0.5 objective lens (Zeiss). The fluorescence intensity of the green (SYTO® 9) and red (propidium iodide) channels was quantified using the FIJI distribution of ImageJ (https://imagej.net/ij/index.html). The values were converted into percentages, with the sum of both channels totaling 100%, allowing comparison of the proportion of live and dead cells between 0 and 30 min of sanitization for each surface and treatment. All images were processed using the same software previously mentioned (FIJI), exclusively for channel separation and overlay, as well as for brightness and contrast adjustments.

### 2.7 Statistical analysis

Statistical analysis of the sanitizer performances was conducted based on the microbial load reduction values (log N – log N_₀_). According to the Shapiro–Wilk test, the dataset showed a normal distribution. The data was analyzed by ANOVA, and Bonferroni correction was applied for multiple comparisons between group interactions. Analyses were performed using the R statistical environment (R Core Team, 2024).

## 3. Results

### 3.1 DSB quantification after dry sanitization

DSB counts for *Salmonella* spp. on SS and PP surfaces were performed before and after each sanitizer exposure time (Figure 2). Overall, the DSB formation endpoint count was around 7.0 log CFU/cm² for P1 (48-h hydrated + 48-h dry) and between 5.0 and 7.0 log CFU/cm² for P2 (24-h hydrated + 120-h dry). In contrast, P3 (8-h hydrated + 48-h dry) resulted in a less dense biofilm, with counts ranging from 3.0 to 6.0 log CFU/cm².

**Figure 2.**
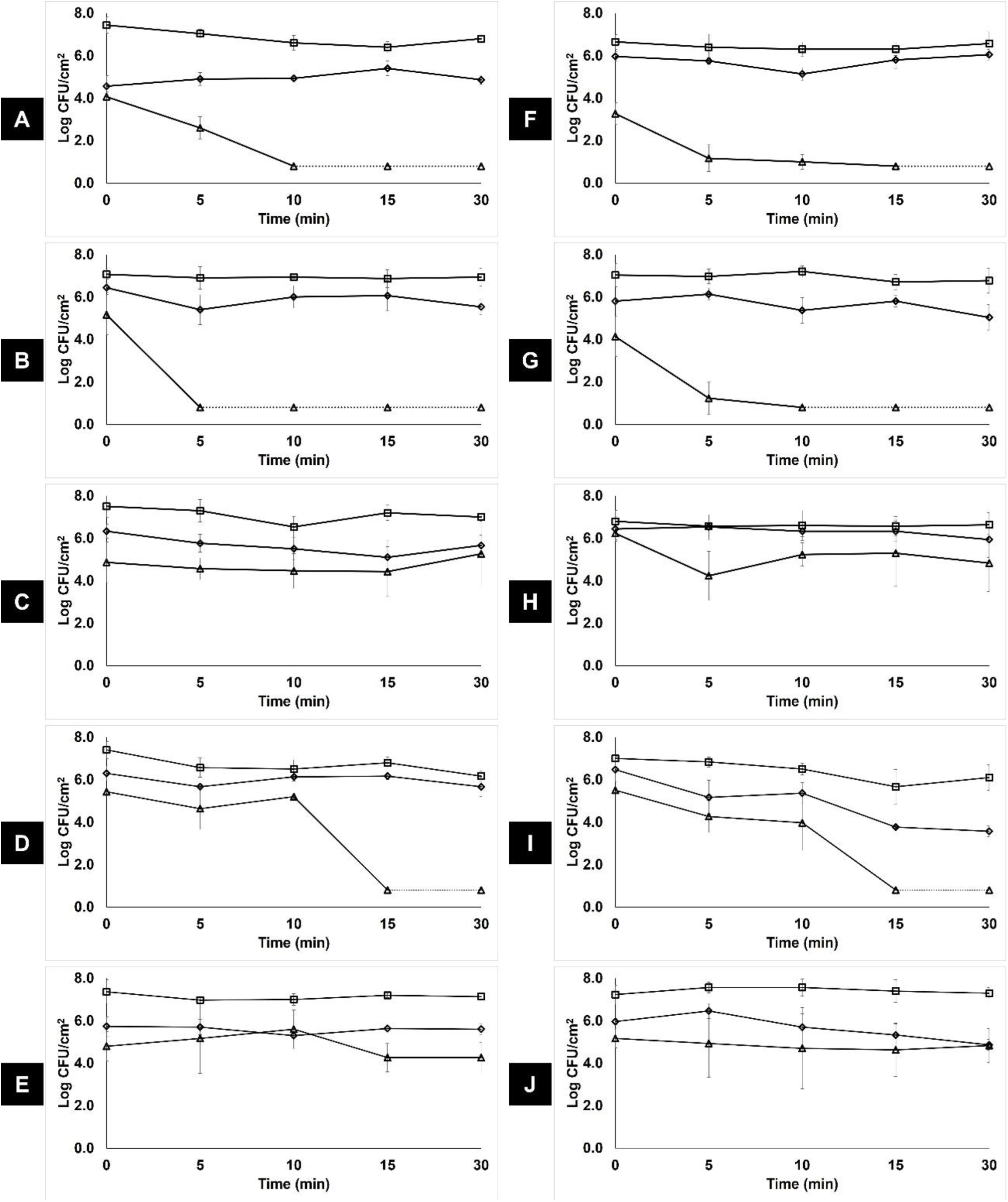
Adherent cell counts (AC) of Salmonella dry surface biofilm on P1 - ▪, P2 - ◆, and P3 - ▴ on surfaces of 1) Stainless Steel – SS (A - E) and Polypropylene – PP (F - J) after sanitization with: (A, F) 70% alcohol - 70%, (B, G) product containing QAC - 0.015% and IPA - 25%, (C, H) hot air - 90°C, (D, I) UV-C irradiation - 254 nm, (E, J) gaseous ozone - 45 ppm.

The results show that treatment time, the type of DSB protocol, and the surface material had a heterogeneous effect on the efficacy of the sanitizers. For 70% ethanol, the greatest reductions were observed in P3 (p < 0.05), with counts below the detection limit (0.8 log CFU/cm²) after 10 min on SS and 15 min on PP. Meanwhile, for DSB formed with P1, reductions of up to 0.4 and 1.0 log CFU/cm² were recorded on PP and SS, respectively. For DSB formed with P2, the greatest reduction obtained in PP was 0.8 log CFU/cm² after 10 min of treatment, whereas in SS no reduction was observed over the 30 min of exposure to the sanitizer (Figure 2a and 2f). There was a significant difference in DSB counts following 70% ethanol exposure (p < 0.05) between P1 and P2 only in SS.

The commercial product (Figures 2b and 2g) exhibited behavior similar to that of 70% ethanol. However, there was a significant difference (p < 0.05) between the types of surfaces and the DSB protocols. Only P3 reached the detection limit after 5 min on SS and 10 min on PP (p < 0.05). Following P1 DSB formation, counts ranged from 6.9 to 7.1 log CFU/cm² on SS and from 6.7 to 7.2 log CFU/cm² on PP (p < 0.05). For DSB formed with P2, counts ranged from 5.4 to 6.4 log CFU/cm² in SS and from 5.0 to 6.1 log CFU/cm² in PP (p > 0.05).

Following hot-air treatment (Figures 2c and 2h), reduction in bacterial count did not reach the limit of detection after 30 minutes of exposure regardless of the DSB formation protocol. Following the DSB formation protocols P1 and P2, bacterial reduction observed were ≤1.0 and ≤1.2 log CFU/cm² in SS and ≤0.2 and ≤0.5 log CFU/cm² in PP, respectively. For DSB formed with P3, the greatest reduction was obtained on PP after 5 min (2.0 log CFU/cm²), while on SS the reductions were ≤0.4 log CFU/cm², although there was no significant difference in bacterial reduction between the surfaces (p > 0.05).

Exposure to UV-C light for 30 minutes resulted in reductions of 1.2 log CFU/cm² on SS and 0.9 log CFU/cm² on PP when DSB were formed following P1. With DSB formed with P2, reductions of 0.6 log CFU/cm² on SS and 2.9 log CFU/cm² on PP were observed. Meanwhile, the use of P3 to form DSB, resulted in bacterial counts that reached the detection limit (0.8 log CFU/cm²) after 15 minutes on both surfaces, with reductions > 4.5 log CFU/cm² (Figure 2d and 2i). There was a significant difference in bacterial reduction (p < 0.05) between the surfaces, and between DSB formed with P3 (p < 0.05) and those formed with P1 or P2.

Following treatment with gaseous ozone, bacteria in DSB formed with any of the protocols showed slight reductions over the course of exposure (≤ 0.5 log CFU/cm²), with the exception of DSB form on PP following P2, where a reduction of 1.1 log CFU/cm² was observed after 30 min. There was no significant difference in bacterial count between the surfaces (p > 0.05). However, there was a difference in bacterial recovered from PP (p < 0.05) between DSB produced with P1 and P3 after 15 min of exposure, and between P1 and P2 and P3 after 30 min.

### 3.2 CLSM Analysis

CLSM analysis was used to assess the cell viability of *Salmonella* spp. DSB before (forming endpoint, 0 min) and after 30 min of exposure to sanitizers. Before sanitization, DSB exhibited predominantly green fluorescence on both surfaces, indicating a high proportion of viable cells (Figure 3). Nevertheless, the images revealed heterogeneity among samples within the same protocol, between DSB protocols, and across surface types, highlighting the challenges associated with assay reproducibility. In P1 and P3, SS exhibited a higher proportion of red-stained (dead cells) than PP. Overall, CLSM images revealed that, even after treatment, the biofilms, in most of the cases, remained present and viable on the surfaces and exhibited a high percentage of green fluorescence (Figure 3), with the exception of DSB on SS formed with P1. The presence of yellow-stained cells, observed in some treatments, may indicate partial membrane damage with penetration of both dyes into the cell cytoplasm.

**Figure 3.**
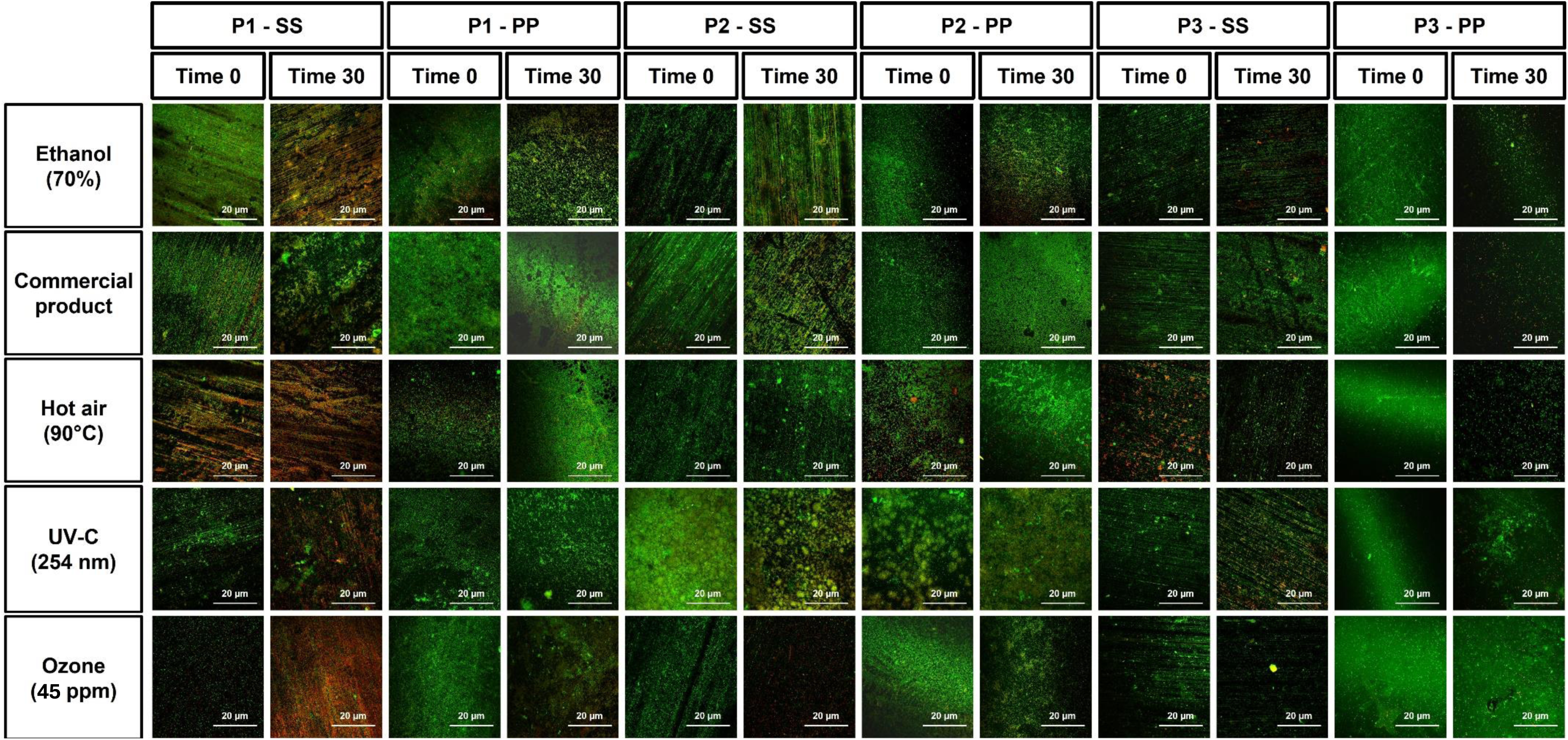
Confocal laser microscopy (live/dead) of three Salmonella dry-surface biofilm protocols (P1 (48h/48h), P2 (24 h/120 h) and P3 (8 h/48 h)) formed on stainless steel (SS) and polypropylene (PP) coupons and subjected to sanitization for 0 and 30 min.

## 4. Discussion

DSB initial counts observed in the current study following DSB formation protocols P1 and P2, corroborate with Vaz et al. (2026) who reported counts between 6.0 and 7.0 log CFU/cm² for *Salmonella* DSB. In contrast, bacteria count from DSB following P3 resulted in a low cell count, being significantly different (p < 0.05) from other protocols. Vaz et al. (2026) also reported a less dense biofilm using a protocol similar to P3.

There is no official methodology in the U.S. or Europe for evaluating the efficacy of sanitizers classified as dry methods against DSB. Our results showed that the type of formation protocol significantly influenced the performance of sanitizers (p < 0.05). There was limited activity against denser DSB formed following successive equal hydrated and desiccation phases (P1) or those that had undergone prolonged desiccative stress (P2; 120 h dry phase). In contrast, the DSB protocol with short hydrated phase humidity (P3, 8-h/48-h) showed greater susceptibility. Three of the five sanitizers evaluated resulted in significant reductions in bacteria in DSB, with counts below the detection limit after 5 to 15 min of exposure (Figure 2). This would suggest that the duration of the hydrated phase plays a relevant role in *Salmonella* DSB formation and, consequently, in sanitizer resistance. A lower cell concentration, and possibly a lower EPS concentration were characterized as a scenario more favorable to the penetration and action of sanitizers.

When comparing the performance of the sanitizers against DSB formed on the same surface using the same DSB formation protocol, it was observed that for DSB formed following P1 and P2, UV-C differed from the other treatments starting at 15 min exposure, with reductions of up to 2.9 log CFU/cm² after 30 minutes. Although UV-C showed the best performance among the method evaluated, its limited action can be explained by its interaction with organic matter from EPS and its low penetration power in dense biofilms, such as DSB (Park et al. 2024). However, in *Salmonella* hydrated surface biofilm (HSB), von Hertwig et al. (2023) reported reductions of up to 4.2 log on PP and 2.2 log on SS after 30 min of exposure to UV-C.

On the other hand, following DSB formation with P3, 70% alcohol, commercial product, and UV-C differed from hot air and ozone after 5 to 10 minutes of exposure. Alcohol-based sanitizers are widely used in the low-moisture industry. As reported by Lin et al. (2024), DSB cells can be surrounded by a dense capsule that protects them against sanitizer action. Alcohol-based sanitizers can precipitate the exopolysaccharides that compose this capsule, disrupting its structure and enabling sanitizer penetration. It is likely that DSB formed with short exposure to wet condition are more susceptible to damage, as their capsule layer is less developed, which is probably what occurred in DSB formed with P3 where hydration phase was shorter (Figure 2). Boyce (2018) indicates that organic matter, such as EPS, can reduce the effectiveness of these sanitizers, which could explain the low performance observed against DSB formed with P1 and P2, with reductions < 1.0 log CFU/cm^2^. Nevertheless, it is important to note that EPS concentration was not evaluated in the current study. Lin et al. (2024) also reported a limited action of isopropyl on *Salmonella* DSB formed in a Petri dish, with reduction of up to 1.6 log CFU/sample.

In contrast, hot air and ozone performed poorly against DSB formed with any of the protocols investigated. The low efficiency of hot air may be associated with heat dissipation and the difficulty of achieving lethal temperatures uniformly across the entire surface of the biofilm, particularly on PP, which acts as a thermal insulator (Astrouski et al, 2020; Harada and Nascimento, 2021b). In addition, desiccation stress induces synthesis of heat-shock proteins and chaperones (Gruzdev et al., 2012; Wang et al., 2021). However, hot air had the best performance on *Salmonella* HSB formed on SS compared to our results, with reductions up to 3.3 log after 30 min exposure (von Hertwig et al., 2023).

Finally, the limited effectiveness of ozone may be related to its reactivity, which restricts its penetration into the polymeric matrix of the biofilm (Viera et al., 1999; Marino et al, 2018). The low penetration of ozone into dense biofilms is well documented and may explain the absence of significant reductions (Panebianco et al., 2021). The superoxide dismutase enzyme has an antioxidant capacity, and its action may be related to antimicrobial resistance to ozone (Sun et al., 2022).

There was a significance impact (P < 0.05) of surface type on the efficacy of the commercial product (QAC + isopropyl alcohol) against DSB formed with P1 and P3, and of UV-C light against DSB formed with P2 and P3, indicating that the nature of the surface influences the performance of the sanitizers. Siuh observation be explained due to the poor penetration capability of UV-C and the shadowing effect caused by surface irregularities (Gabriel et al., 2018; Turtoi, 2013). In addition, UV-C can cause photodegradation in PP, resulting in photo-oxidation and a decrease in the hardness of the material contributing to the microbial inactivation (Kowalski, 2009).

In general, CLSM images corroborate with plate counts (log CFU/cm²). However, some discrepancies were observed. The P3 population treated with the commercial product was rapidly reduced to counts below the detection limit, as mentioned earlier. Nevertheless, under the microscope, the samples showed a high proportion of cells stained green (SYTO 9), which could be attributed to the presence of viable but nonculturable (VBNC) and sublethal injured cells (Ramamurthy et al., 2014). Under these conditions, the cells can maintain reduced metabolic activity and membrane integrity, although they lose their ability to proliferate in conventional culture media (Centeleghe et al, 2023; Jean-Marie et al, 2025). Lin et al. (2024) and Vaz et al. (2026) previously demonstrated the presence of VBNC cells in *Salmonella* DSB. In addition, these results may reflect inherent limitations of membrane-integrity-based dyes, which can lead to an overestimation of cell viability (Berney et al., 2007; Ramamurthy et al., 2014, Luo and Raval, 2025). On the other hand, a high percentage of red-stained cells in treatments that exhibited high colony counts, as observed for gaseous ozone, may indicate the occurrence of cellular stress associated with the formation of double-strand breaks (DSB) and exposure to disinfectants. Under these conditions, a transient increase in cell membrane permeability may occur, allowing the entry of propidium iodide without irreversibly compromising viability and culture capacity (Berney et al., 2007; Davey, 2011). This phenomenon could also characterize a sublethal effect of the sanitizer in question.

Our results indicate that the efficacy of the dry sanitization methods evaluated in this study is impacted by variables such as exposure time, the surface material and in particular the duration of exposure to high humidity during DSB formation. In fact, P3 was the easiest DSB to be controlled. However, it is worth noting that no treatment achieved reductions > 3 logs, indicating limited efficacy of the sanitizers. DSB resistance to disinfection compared to HSB has been reported (Centeleghe et al., 2023, Lin et al., 2024). When *Salmonella* DSB are considered, the same conclusion could be drawn, notably when comparing our results to those reported by von Hertwig et al. (2023), who challenged *Salmonella* HSB against dry sanitizers and obtained bacterial reductions ranging from 1.3 to 4.2 log CFU/cm². These results can be related with dense biofilm and modifications in the cell membrane caused by the desiccation stress (Lin et al., 2024; Maillard & Centeleghe, 2023; Rahman et al., 2022). Additional studies on the combination of physical and chemical methods are needed to optimize sanitation processes in environments with controlled humidity, such as low-moisture food industries.

## CRediT authorship contribution statement

**Vinícius S. A. Vaz:** Writing – original draft, Visualization, Investigation, Formal analysis, Data curation. **Jéssica de A.F.F. Finger:** Writing – original draft, Visualization, Formal analysis, Data curation. **Raul F. Pereira:** Methodology, Investigation, Formal analysis. **Erika S. Silva:** Methodology, Investigation, Formal analysis. **Rafael P. Maia:** Investigation, Statistics analysis. **Jean-Yves Maillard:** Methodology, Writing – review & editing**. Maristela S. Nascimento:** Writing – review & editing, Resources, Project administration, Methodology, Funding acquisition, Conceptualization, Data curation.

## Acknowledgements

The authors acknowledge financial support from Fundação de Amparo à Pesquisa do Estado de São Paulo (FAPESP; 2021/06809-2 and 2023/03076-0), Conselho Nacional de Desenvolvimento Científico e Tecnológico (CNPq; 305702/2021-1), and Coordenação de Aperfeiçoamento de Pessoal de Nível Superior – Brazil (CAPES; Finance Code 001). The authors also thank the National Institute of Science and Technology on Photonics Applied to Cell Biology (INFABIC) at the State University of Campinas for providing access to CLSM equipment and technical support (FAPESP; 2014/50938-8 and CNPq; 465699/2014-6).

## References

Alonso, V. P. P., Gonçalves, M. P. M. B. B., de Brito, F. A. E., Barboza, G. R., Rocha, L. de O., Silva, N. C. C., (2023). Dry surface biofilms in the food processing industry: An overview on surface characteristics, adhesion and biofilm formation, detection of biofilms, and dry sanitization methods. Comprehensive Reviews in Food Science and Food Safety, 22(1), 688–713. 10.1111/1541-4337.13089.

Andrade, N.J.D., (2008). Higienização na indústria de alimentos: avaliação e controle da adesão e formação de biofilmes bacterianos. Viçosa: Editora UFV, 412 pp.

Astrouski, I., Raudensky, M., Kudelova, T., & Kroulikova, T., (2020). Fouling of Polymeric Hollow Fiber Heat Exchangers by Air Dust. Materials, 13, Article 4931. 10.3390/ma13214931

Baker, J., Rana, Y. S., Chen, L., Beary, M. A., Balasubramaniam, V. M., Snyder, A. B., (2025). Superheated steam can rapidly inactivate bacteria, but manual operation of commercial units resulted in limited efficacy during dry surface sanitization. Journal of Food Protection, 88, Article 100461. 10.1016/j.jfp.2025.100461.

Berney, M., Hammes, F., Bosshard, F., Weilenmann, H.U., Egli, T., (2007). Assessment and Interpretation of Bacterial Viability by Using the LIVE/DEAD BacLight Kit in Combination with Flow Cytometry. Applied Environmental Microbiology, 73(10), 3283–3290. 10.1128/AEM.02750-06.

Boyce, J. M., (2018). Alcohols as surface disinfectants in healthcare settings. Infection Control and Hospital Epidemiology, 39(3), 323–328. 10.1017/ice.2017.301.

Centeleghe, I., Norville, P., Hughes, L., & Maillard, P., (2023). Klebsiella pneumoniae survives on surfaces as a dry biofilm. American journal of infection control, 51, 1157–1162. 10.1016/j.ajic.2023.02.009.

Centers for Disease Control and Prevention (CDC), (2025). *Salmonella* Infection (Salmonellosis). About Salmonella Infection. Retrieved from https://www.cdc.gov/salmonella/about/index.html. Accessed August 8, 2025.

Codex Alimentarius, (2018). Code of Hygienic Practice for Low-Moisture Foods. CXC 75-2015. Retrieved from https://www.fao.org/fao-who-codexalimentarius/sh-proxy/fr/?lnk=1&url=https%253A%252F%252Fworkspace.fao.org%252Fsites%252Fcodex%252FStandards%252FCXC%2B75-2015%252FCXC_075e.pdf. Accessed August 12, 2025.

Davey, H.M., (2011). Life, Death, and In-Between: Meanings and Methods in Microbiology. Applied Environmental Microbiology, 77(16), 5571–5576. 10.1128/AEM.00744-11.

Food and Agriculture Organization of the United Nations and World Health Organization, (2022). Ranking of low moisture foods in support of microbiological risk management: meeting report and systematic review. Microbiological Risk Assessment Series No. 26. Rome. 10.4060/cc0763en.

Gabriel, A. A., Ballesteros, M. L. P., Rosario, L. M. D., Tumlos, R. B., Ramos, H. J., (2018). Elimination of Salmonella enterica on common stainless steel food contact surfaces using UV-C and atmospheric pressure plasma jet. Food Control, 86, 90–100. 10.1016/J.FOODCONT.2017.11.011.

Galié, S., García-Gutiérrez, C., Miguélez, E.M., Villar, C.J., Lombó, F., (2018) Biofilms in the Food Industry: Health Aspects and Control Methods. Frontiers in Microbiology. 9, 315815. 10.3389/fmicb.2018.00898.

Gruzdev, N., Pinto, R., Sela Saldinger, S., (2012). Persistence of Salmonella enterica during dehydration and subsequent cold storage. Food Microbiology, 32, 415–422. 10.1016/j.fm.2012.08.003.

Harada, A. M. M., Nascimento, M. S., (2021a). Effect of dry sanitizing methods on Bacillus cereus biofilm. Brazilian Journal of Microbiology, 52(2), 919–926. 10.1007/s42770-021-00451-0.

Harada, A. M. M., Nascimento, M. S., (2021b). Efficacy of dry sanitizing methods on Listeria monocytogenes biofilms. Food Control, 124, Article 107897. 10.1016/j.foodcont.2021.107897.

Ivers, C., Chalamalasetti, S., Ruiz-Llacsahuanga, B., Critzer, F., Bhullar, M., Nwadike, L., Yucel, U., Trinetta, V., (2024). Evaluation of commercially available sanitizers efficacy to control Salmonella (sessile and biofilm forms) on harvesting bins and picking bags. Journal of Food Protection, 87, Article 100394. 10.1016/J.JFP.2024.100394.

Jean-Marie, N., Lebielle, T., Louisin, M., Olive, C., Marion-Sanchez, K., (2025). A fully automated model to form "dry surface biofilms" under optimal dehydration conditions. application to Enterobacteriaceae in healthcare settings. Biofilm, 10, Article 100312. 10.1016/j.bioflm.2025.100312.

Karygianni, L., Ren, Z., Koo, H., Thurnheer, T., (2020). Biofilm matrixome: extracellular components in structured microbial communities. Trends in Microbiology. 28, 668–681. 10.1016/j.tim.2020.03.016.

Kowalski, W., (2009). Ultraviolet germicidal irradiation handbook: UVGI for air and surface disinfection. Springer science & business media. 10.1007/978-3-642-01999-9.

Ledwoch, K., Said, J., Norville, P., Maillard, J. Y., (2019). Artificial dry surface biofilm models for testing the efficacy of cleaning and disinfection. Letters in Applied Microbiology, 68, 329–336. 10.1111/lam.13143.

Ledwoch, K., Vickery, K., Maillard, J. Y., (2022). Dry surface biofilms: what you need to know. British Journal of Hospital Medicine, 83, 1–3. 10.12968/hmed.2022.0274.

Lin, Z., Liang, Z., He, S., Chin, F. W. L., Huang, D., Hong, Y., Wang, X., & Li, D., (2024). *Salmonella* dry surface biofilm: morphology, single-cell landscape, and sanitization. Applied and Environmental Microbiology, 90, Article e01623-24. 10.1128/aem.01623-24.

Liu, S., Roopesh, M. S., Tang, J., Wu, Q., Qin, W., (2022). Recent development in low-moisture foods: Microbial safety and thermal process. Food research international, 155, Article 111072. 10.1016/j.foodres.2022.111072.

Luo, J., Raval, R., (2025). Correlative Imaging and super resolution microscopy studies reveal complexities in determining live-dead state of bacteria. Biofilm, 10, Article 100302. 10.1016/j.bioflm.2025.100302.

Maillard, J. Y., Centeleghe, I., (2023). How biofilm changes our understanding of cleaning and disinfection. Antimicrobial Resistance & Infection Control, 12, Article 95. 10.1186/s13756-023-01290-4

Marino, M., Maifreni, M., Baggio, A., Innocente, N., (2018). Inactivation of Foodborne Bacteria Biofilms by Aqueous and Gaseous Ozone. Frontiers in Microbiology. 28, 2024. 10.3389/fmicb.2018.02024

Milton, A. A. P., Srinivas, K., Lyngdoh, V., Momin, A. G., Lapang, N., Priya, G. B., Ghatak, S., Sanjukta, R. K., Sen, A., Das, S., (2023). Biofilm-forming antimicrobial-resistant pathogenic *Escherichia coli*: A one health challenge in Northeast India. Heliyon, 9, Article e20059. 10.1016/j.heliyon.2023.e20059.

Moerman, F., Mager, K., (2016). Cleaning and disinfection in dry food processing facilities. *In* Handbook of hygiene control in the food industry (pp. 521-554). Woodhead Publishing. 10.1016/B978-0-08-100155-4.00035-2.

Nascimento, M. S., Carminati, J. A., Silva, I. C. R. N., Silva, D. L., Bernardi, A. O., Copetti, M. V., (2018). *Salmonella*, *Escherichia coli* and *Enterobacteriaceae* in the peanut supply chain: from farm to table. Food Research International, 105, 930–935. 10.1016/j.foodres.2017.12.021.

Nerney, A., Reitz, S., Kovacevic, J., Waite-Cusic, J., (2025). Cross-contamination risks in dry produce packinghouses: efficacy of alcohol-based sanitizers to reduce salmonella and potential surrogates on relevant surface materials. Journal of Food Protection, 88, Article 100443. 10.1016/j.jfp.2024.100443.

Nicholas, R., Dunton, P., Tatham, A., Fielding, L., (2013). The effect of ozone and open air factor on surface-attached and biofilm environmental Listeria monocytogenes. Journal of Applied Microbiology, 115, 555–564. 10.1111/jam.12239.

Olanbiwoninu, A. A., Popoola, B. M., (2023). Biofilms and their impact on the food industry. Saudi Journal of Biological Sciences, 30, Article 103523. 10.1016/J.SJBS.2022.103523.

Otter, J. A., Vickery, K., Walker, J. D., Pulcini, E. D., Stoodley, P., Goldenberg, S. D., Salkeld, J.A.G., Chewins, J., Yezli, S., Edgeworth, J. D., (2015). Surface-attached cells, biofilms and biocide susceptibility: implications for hospital cleaning and disinfection. Journal of Hospital Infection, 89, 16–27. 10.1016/j.jhin.2014.09.008.

Otto, C., Zahn, S., Rost, F., Zahn, P., Jaros, D., Rohm, H., (2011). Physical methods for cleaning and disinfection of surfaces. Food Engineering Reviews, 3, 171–188. 10.1007/s12393-011-9038-4.

Panebianco, F., Rubiola, S., Chiesa, F., Civera, T., Di Ciccio, P. D., (2021). Effect of gaseous ozone on *Listeria monocytogenes* planktonic cells and biofilm: an in vitro study. Foods, 10. 10.3390/foods10071484

Park, H. W., Balasubramaniam, V. M., Snyder, A. B., (2024). Inactivation of *Enterococcus faecium* and *Geobacillus stearothermophilus* spores on stainless steel through dry sanitation approaches using superheated steam and ultraviolet C-LED. Food Control, 156, Article 110144. 10.1016/j.foodcont.2023.110144

Prestes, F. S., Yotsuyanagi, S. E., Alonso, V. P., Nascimento, M. S., (2024). Dry sanitization in the food industry: a review. Current Opinion in Food Science, 57, Article 101166. 10.1016/J.COFS.2024.101166.

R Core Team, (2024). *R: A Language and Environment for Statistical Computing.* R Foundation for Statistical Computing, Vienna, Austria. https://www.R-project.org/.

Rahman, M. A., Amirkhani, A., Chowdhury, D., Mempin, M., Molloy, M. P., Deva, A. K., Vickery, K., & Hu, H., (2022). Proteome of *Staphylococcus aureus* Biofilm Changes Significantly with Aging. International journal of molecular sciences, 23, Article 6415. 10.3390/ijms23126415.

Ramamurthy, T., Ghosh, A., Pazhani, G.P., Shinoda, S., (2014). Current Perspectives on Viable but Non-Culturable (VBNC) Pathogenic Bacteria. Frontiers Public Health, 2, 103. 10.3389/fpubh.2014.00103.

Rosado, M. S., (2009). Biofilme de *Enterococcus faecium* em superfície de aço inoxidável: caracterização tecnológica, modelagem e controle por agentes sanitizantes. Dissertação (Mestre em Tecnologia de Alimentos) - Faculdade de Engenharia de Alimentos, Universidade Estadual de Campinas, Campinas (SP), 84p.

Sadiq, F. A., De Reu, K., Burmølle, M., Maes, S., Heyndrickx, M., (2023). Synergistic interactions in multispecies biofilm combinations of bacterial isolates recovered from diverse food processing industries. Frontiers Microbiology, 14, Article 1159434. 10.3389/fmicb.2023.1159434.

Sauer, K., Stoodley, P., Goeres, D. M., Hall-Stoodley, L., Burmølle, M., Stewart, P. S., Bjarnsholt, T., (2022). The biofilm life cycle: expanding the conceptual model of biofilm formation. Nature Reviews Microbiology. 20, 608–620. 10.1038/s41579-022-00767-0.

Sun, Y., Li, J., Yang, Y., Yang, G., Shi, Y., Wang, S., Wang, M., Xia, X., (2022). The Role of ptsH in Stress Adaptation and Virulence in Cronobacter sakazakii BAA-894. Foods, 11, Article 2680. 10.3390/foods11172680.

Turtoi, M., (2013). Ultraviolet light treatment of fresh fruits and vegetables surface: A review. Journal of Agroalimentary Processes and Technologies, 19.3, 325–337.

Vaz, V. S. A., de A.F.F. Finger, J., Pereira, R. F., Derami, M. S., Maillard, J. Y., Nascimento, M. S., (2026). Dry surface biofilm of Salmonella and Cronobacter sakazakii: a real concern for the low moisture food industry. Food Microbiology, 136, Article 105013. 10.1016/J.FM.2025.105013.

Viera, M. R., Guiamet, P. S., De Mele, M. F. L., Videla, H. A., (1999). Biocidal action of ozone against planktonic and sessile *Pseudomonas fluorescens*. Biofouling, 14, 131–141. 10.1080/08927019909378404

von Hertwig, A. M, Prestes, F. S., do Nascimento, M. D. S., (2022). Biofilm formation and resistance to sanitizers by Salmonella spp. Isolated from the peanut supply chain. Food Research International, 152, Article 110882. 10.1016/j.ijfoodmicro.2019.02.005.

von Hertwig, A. M., Prestes, F. S., Nascimento, M. S., (2023). Comparative evaluation of the effectiveness of alcohol-based sanitizers, UV-C radiation and hot air on three-age Salmonella biofilms. Food Microbiology, 113, Article 104278. 10.1016/J.FM.2023.104278.

Wang, L., Forsythe, S. J., Yang, X., Fu, S., Man, C., Jiang, Y., (2021). Invited review: Stress resistance of Cronobacter spp. affecting control of its growth during food production. Journal of Dairy Science, 104, 11348–11367. 10.3168/jds.2021-20591.

Wang, Y., Feng, Y., Wang, X., Ji, C., Upadhyay, A., Xiao, Z., Luo, Y., (2025). A short review on *Salmonella* spp. involved mixed-species biofilm on food processing surface: interactions, disinfectant resistance and its biocontrol. *Journal of Agriculture and Food Research*, Article 101660. 10.1016/j.jafr.2025.101660.

Zhong, W., Alfa, M., Zelenitsky, S., Howie, R., (2009). Simulation of cyclic reprocessing buildup on reused medical devices. Computers in Biology and Medicine, 39, 568–577. doi:10.1016/j.compbiomed.2009.04.003

